# Structures of perforin-2 in solution and on a membrane reveal mechanisms for pore formation

**DOI:** 10.1101/2022.06.14.496043

**Authors:** Xiulian Yu, Tao Ni, George Munson, Peijun Zhang, Robert J. C. Gilbert

## Abstract

Perforin-2 (PFN2, MPEG1) is a key pore-forming protein in mammalian innate immunity restricting intracellular bacteria proliferation. It forms a membrane-bound pre-pore complex that converts to a pore-forming structure upon acidification; but its mechanism of conformational transition has been debated. Here we used cryo-electron microscopy, tomography and subtomogram averaging to determine structures of PFN2 in pre-pore and pore conformations in isolation and bound to liposomes. In isolation and upon acidification, the pre-assembled complete pre-pore rings convert to pores in both flat ring and twisted conformations. The twisted pore structure suggests an intermediate or alternative state to the flat conformation, and a capacity to distort the underlying membrane during membrane insertion. On membranes, *in situ* assembled PFN2 pre-pores display various degrees of completeness; whereas PFN2 pores are mainly incomplete arc structures that follow the same subunit packing arrangements as found in isolation. Both assemblies on membranes use their P2 β-hairpin for binding to the lipid membrane surface. These structural snapshots in different states reveal a molecular mechanism for PFN2 pre-pore to pore transition on a targeted membrane.

## Introduction

Innate immunity plays key roles in defense against invading pathogens such as bacteria and viruses. In mammals, several families of pore forming proteins are involved in this process, including lymphocytic perforin-1, the complement system membrane attack complex (MAC), and the recently-identified perforin-2 (hereafter PFN2, (McCormack *et al*, 2013; Merselis *et al*, 2021)), which is evolutionarily the most ancient pore forming protein in immune defense (McCormack & Podack, 2015). PFN2 plays a pivotal role in phagocytes for the restriction of intracellular bacteria, and is thought to form pores in targeted bacterial membranes: human individuals carrying PFN2 missense mutations are significantly more susceptible to bacterial infection (McCormack *et al*, 2017; Merselis *et al*, 2020); and PFN2 knockout mice have bacterial pathogens replicating and disseminating more deeply into their tissues (McCormack *et al*, 2016; McCormack & Podack, 2015; McCormack *et al*, 2015b). Besides bacterial restriction, PFN2 is also involved in the release of IL-33 from conventional dendritic cells through their cell membranes, potentially by making pores in the cell membrane that allow IL-33 to transit through (Hung *et al*, 2020).

PFN2 belongs to the membrane attack complex-perforin (MACPF) family and shares structural homology with bacterial cholesterol-dependent cytolysins (CDCs) (Gilbert *et al*, 2013). The mechanism of pore formation has been studied extensively for many members of the MACPF/CDC superfamily, and usually involves the formation of a pre-pore structure on a target membrane (such as a macrophage or bacterium) which transitions to a pore-forming state (Gilbert, 2014). A large conformational change occurs during pre-pore to pore transition, including domain reorganization and refolding such that two sets of amphipathic α-helices (TMH1 and TMH2) in the MACPF/CDC domain in the pre-pore conformation convert to a 4-stranded β-sheet (Gilbert, 2014; Shepard *et al*, 2000; Tilley *et al*, 2005). PFN2 is expected to function in a similar way.

Distinct from MAC and perforin-1, PFN2 is a type I transmembrane protein that is expected to be cleaved from its transmembrane anchor during bacterial restriction, releasing from the host membrane an ectodomain consisting of the MACPF and membrane-binding P2 domain (McCormack & Podack, 2015; McCormack *et al*, 2015a). Previous work revealed structures for pre-pore and pore states of murine PFN2 (mPFN2) assembled in solution (hereafter referred as isolated assembly vs membrane-bound assembly), showing that mPFN2 undergoes substantial conformational rearrangement when transiting from the pre-pore to pore upon acidification (Ni *et al*, 2020). In the pre-pore state the membrane-binding P2 domain is oriented in the opposite direction to the pore-forming MACPF domain, compared with the pore state structure; we therefore proposed that pore formation required the MACPF domain to rotate 180° prior to refolding and membrane insertion (Ni *et al*., 2020). These findings were supported by complementary imaging of pre-pores and pores on membranes using negative-staining electron and atomic force microscopy, and by studies using reversibly locked disulfide mutants. An alternative mechanism of pre-pore to pore transition has also been suggested: work on the human PFN2 pre-pore led to the proposition that hPFN2 could use its P2 β-hairpin interacting with one membrane and insert a MACPF pore into another (Bayly-Jones *et al*, 2020; Pang *et al*, 2019), resulting in a pore-forming complex bridging two membranes without rotating the MACPF domain. Although the pore structure we determined argues strongly against this model (Ni *et al*., 2020), structures of PFN2 assemblies formed on targeted membranes (rather than imaged in isolation) are crucial to solve this debate.

In this work, we used cryo-electron microscopy, tomography and subtomogram averaging to determine the structures of mPFN2 in pre-pore and pore conformations in isolation and bound to liposomes. We identify mPFN2 pore structures in isolation in two different forms (flat ring and twisted), suggesting a possible transitional role for the twisted form during membrane pore formation. The membrane-bound mPFN2 pre-pore and pore structures show that both the pre-pore and pore of mPFN2 use their P2 β-hairpins to interact with the target membrane, confirming that the resulting MACPF pore can form in the same membrane to which the P2 domain binds. In addition, the pores are mainly formed by open arcs of mPFN2 subunits, rather than sixteen-subunit assemblies, whether flat or twisted. These results delineate a mechanism for mPFN2 pore formation at a molecular level.

## Results

### PFN2 pre-pore oligomers in solution and on the membrane

The purified mPFN2 ectodomain undergoes self-oligomerization in solution at neutral pH. As revealed in 2D class averages from cryo-EM single particle analysis, mPFN2 forms a series of assemblies with variable numbers of subunits, peaking at 3 subunit oligomers (Fig EV1A and B). To investigate the native state of mPFN2 oligomers on membranes, we assembled pre-pore structures on liposomes prepared from total *E. coli* lipid extract and performed cryo-electron tomography (cryo-ET). mPFN2 oligomerizes into pre-pore complexes on membranes at a neutral pH (Fig 1A). Subtomogram averaging (STA) and classification of these membrane-bound pre-pore complexes reveals a range of assembles comprising arc structures of variable length but the same curvature (Fig 1B). A subclass with C16 symmetry was identified, which revealed a density map at an overall resolution of 6 Å (Fig 1C and Fig EV2A), with a local resolution ranging from 5.2 Å to 9.2 Å (Fig EV2C). In validation of the map quality, secondary structure elements and several glycosylation sites in the MACPF domain (Asn185 and Asn269) were clearly resolved (Fig 1C-D). Molecular dynamics flexible fitting of the pre-pore structure in isolation (PDB: 6SB3) (Ni *et al*., 2020) was performed to derive a snapshot of the membrane-bound pre-pore mPNF2 complex (see Materials and Methods).

**Figure 1.**
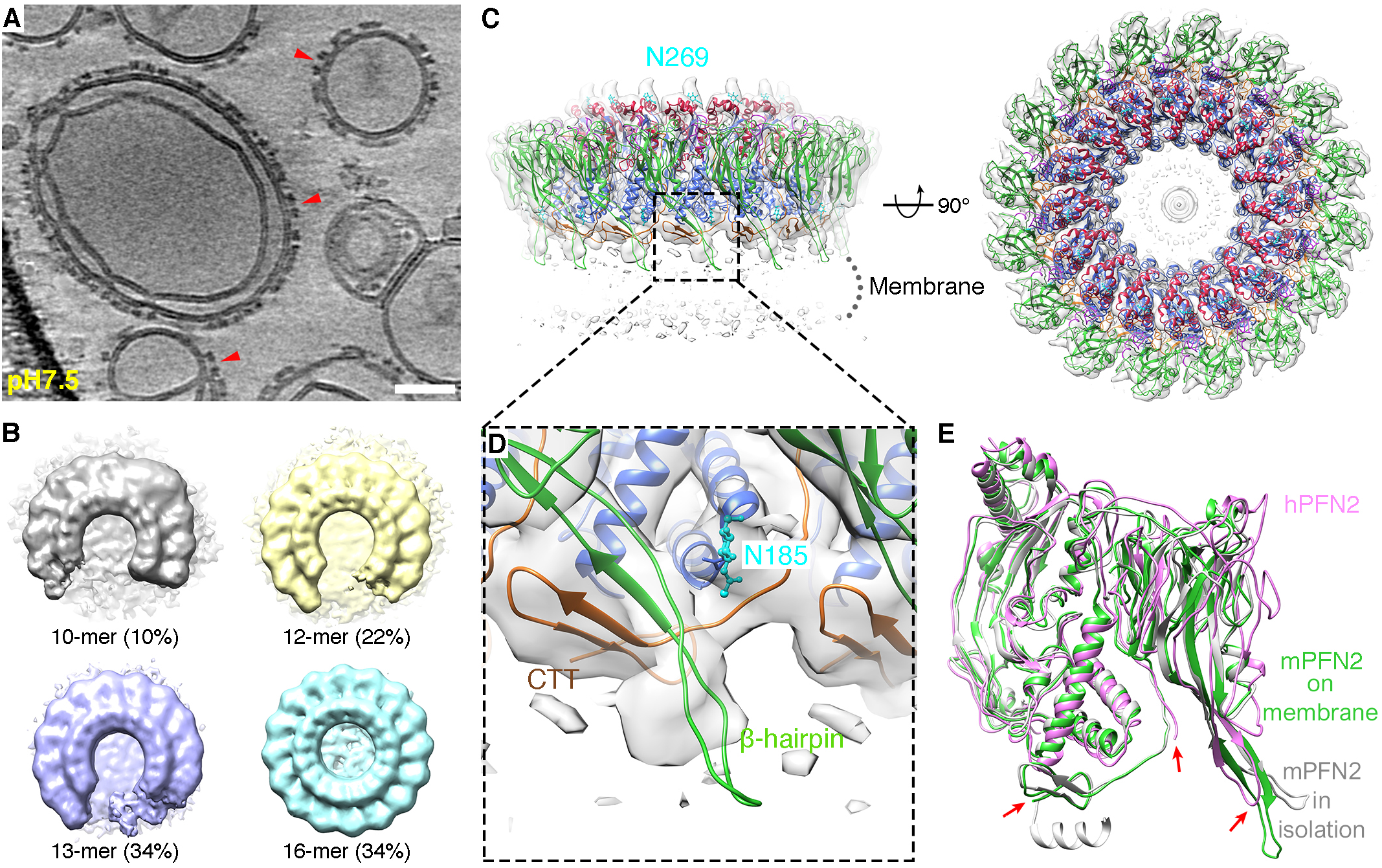
cryoET and subtomogram averaging of mPFN2 pre-pores on liposomes. A. A representative tomogram slice of mPFN2 pre-pores formed on liposomes. Red arrows point to the pre-pores on membrane at side views. Scale bar = 50 nm. Slice thickness 4.68 nm. B. Subtomogram classes of mPFN2 pre-pores assembled on liposomes. C. Structure of mPFN2 pre-pores assembled on liposomes at 6.0 Å resolution by subtomogram averaging, shown in two orthogonal views. Domains are color-coded as MACPF in blue, P2 in green, CTT in orange, the TMH-forming α-helices in red and the conserved N-link glycan in cyan. D. Close-up view of CTT (orange) and β-hairpin (green) of mPFN2 overlaid with density map. E. Comparison of pre-pore PFN2 structures from the human homolog (pink, PDB: 6U2W), mPFN2 in isolation (gray, PDB: 6SB3) and mPFN2 formed on a membrane (green). Red arrows indicate the major differences among the structures.

Overall, the membrane-bound mPFN2 pre-pore structure is similar to that observed in isolation (RMSD: 1.185 Å. Fig 1C-E), except that the tip of the β-hairpin in the P2 membrane binding domain is dissociated from the neighboring subunit and instead extends to the outer membrane leaflet with its hydrophobic tip (Fig 1D-E, green). The C-terminal tail (CTT) is now only partially resolved (residues Ser600 – Lys628, Fig 1D, orange) compared to the in solution pre-pore structure, maintaining an interaction with the P2 β-hairpin from its neighboring subunit (Fig 1D, green) also previously observed (Ni *et al*., 2020). In contrast, the CTT was completely absent in the hPFN2 membrane-bound pre-pore complex determined by cryo-EM SPA (EMD-20622, PDB: 6U2W, Fig 1E, pink). The difference in the CTT structure observed between hPFN2 and mPFN2 on membranes may arise from different approaches to sample preparation: mPFN2 was added as monomers and small oligomers (Fig EV1A-B) and then assembled into pre-pore oligomers after binding to the membrane, while hPFN2 formed double-ring stacked pre-pores spontaneously in isolation during protein purification and was then added to liposomes to obtain a membrane-bound structure (Pang *et al*., 2019). The CTT of the hPFN2 oligomer, which is buried in the middle of the two stacking pre-pores, might have dissociated from the β-hairpin and become disordered when breaking into two independent rings. Indeed, CTT was observed to adopt two different conformations in the isolated ring-stacked hPFN2: it interacts with β-hairpins either from the same subunit or from that of an adjacent subunit, indicating a structural adaptation of this region to its local environment (Pang *et al*., 2019). We previously showed that a CTT truncation mutant can still assemble into the pre-pore oligomer, which suggests that it has an adaptive and possibly regulatory rather than essential role in complex formation, for example in stabilizing the pre-pore state and possibly constituting part of the mPFN2 pH sensor (Ni *et al*., 2020).

The structures of the mPFN2 pre-pore assemblies in solution and on a targeted membrane suggest a dynamic assembly pathway for mPFN2, and reveal that mPFN2 uses the tip of the P2 domain β-hairpin for membrane binding. As in all other pre-pore structures observed (Ni *et al*., 2020; Pang *et al*., 2019), the MACPF domain transmembrane helices are positioned pointing away from the target membrane, while the P2 domain is bound to it.

### Cryo-EM structure of mPFN2 pores identified two major conformations

Having further understood the pre-pore assembly of mPFN2, we next attempted to resolve the structure of mPFN2 pore-forming oligomers both in isolation and on membranes. Our previous structure of the mPFN2 pore at an intermediate resolution revealed its overall architecture as a ring of sixteen subunits (Ni *et al*., 2020). Understanding fully the inter-subunit interactions responsible for the pore assembly, however, requires a higher resolution. As reported previously, pore-forming assemblies were prepared from pre-pore oligomers pre-assembled at pH 5.5, in which condition the bulk of the oligomers are complete full rings (Ni *et al*., 2020), and subsequently by lowering the pH to 3.6-4.0 in the presence of 0.056% Cymal-6 (Ni *et al*., 2020). The isolated converted pores exist in two major conformations, a flat ring conformation (68.5%) and a twisted conformation (15.5%) (Figs 2 and EV3), both structures comprising 16 mPFN2 subunits. The flat ring pore structure was resolved at an overall resolution ∼3.0 Å with C2 symmetry, and with local resolutions ranging from 2.7 to 4.7 Å (Fig EV4). The twisted pore was resolved at ∼4Å with local resolutions ranging from 3.4 to 9.1 Å (Fig EV4).

**Figure 2.**
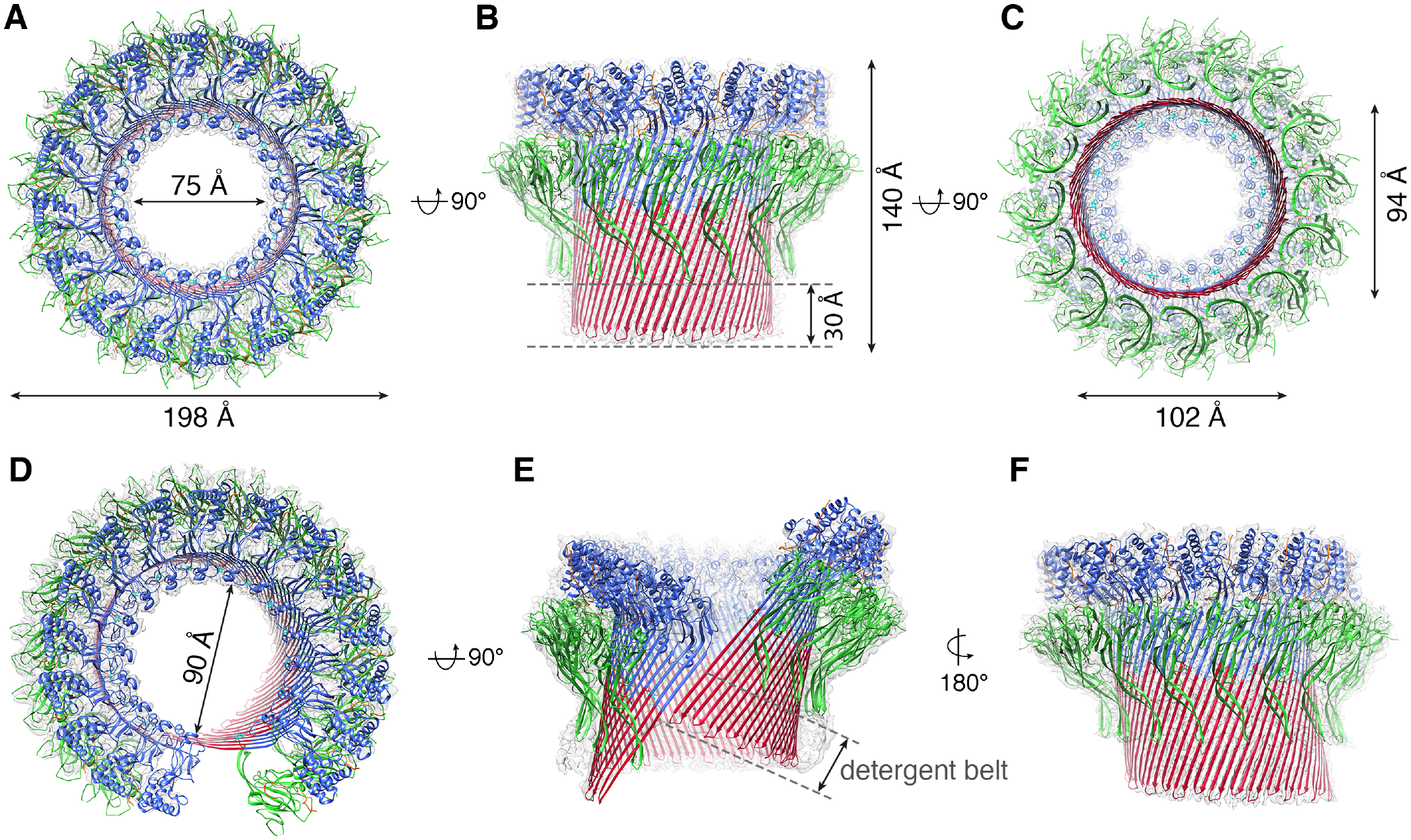
Cryo-EM structures of two mPFN2 pore conformations determined in isolation. A-C, The atomic model of mPFN2 pore-forming structure in a flat ring conformation overlapped with the corresponding cryo-EM map (gray) viewed from top (A), side (B) and bottom (C). D-F, The atomic model of mPFN2 pore-forming structure in a twisted conformation overlapped with the corresponding cryo-EM map (gray) viewed from top (D), front (E) and back (F), the cryoEM map contour level has changed in (E) to show the detergent belt. Domains are color-coded as in Fig1C.

The ring pore follows C16 symmetry in the top of the MACPF domain, but its symmetry is gradually broken towards the bottom of the β-barrel (Fig EV5A). It has an inner diameter of 75 Å at the top of the MACPF domain but a minimum and maximum diameter of 94 Å and 102 Å at the bottom, respectively (Fig 2A-C). 3D variability analysis reveals that the top of the MACPF domain remains largely rigid in organization, presumably due to its large inter-subunit buried surface area, while the bottom of the β-barrel varies between circular and elliptical forms (Fig EV5A and Movie EV1). The twisted pore is slightly larger than the flat ring pore, due partly to a break in the β-strand packing between subunits at the cracking interface, the inner diameter being ∼90 Å (measured between two subunits at the top of MACPF domain, Fig 2D). The sizes of mPFN2 pores indicate that they are sufficiently large to allow the free passage of IL-33 or of agents possibly involved in its antibacterial activity, such a lysozyme, regardless of being in a ring or twisted conformation (Hung *et al*., 2020).

### Inter-subunit interactions within mPFN2 pores

The high-resolution pore structures allow identification of inter-subunit interactions, key to understanding pore assembly. Compared with pre-pore assemblies, several regions became unresolved, including the epidermal growth factor (EGF)-like domain bridging the MACPF and P2 domains, and two β-strands and an α-helix in the very end of the CTT (residues S600 – I642, yellow in the pre-pore, Fig 3A-B). A loop between the P2 domain and CTT, which lies along the MACPF domain in pre-pore oligomers, shifts to interact with its neighboring subunit (residues T581-L590, Fig 3B, red arrow). The rest of the CTT remains tethered to the adjacent MACPF domain, in a similar conformation to that seen in pre-pore structures, although the local density is relatively weak. The flexibility of these regions may facilitate the substantial conformational change of mPFN2 during pore formation, and especially the apparent melting of the EGF-like module’s fold. In addition, the P2 domain rotates ∼90° around its pseudo-3-fold symmetry axis (Fig 3C), and its β-hairpin makes new contacts with the exterior of the β-barrel now presented by the MACPF domain. The β-barrel is further stabilized by a molecule of the detergent Cymal-6, aligned with a strand from the P2 β-hairpin and whose aliphatic region is packed with a series of hydrophobic residues from the back of the TMH barrel in the MACPF domain (Fig 3D). This suggests a role for bilayer components themselves in stabilization of the pore, and a specific acyl chain-protein interface. Surface electrostatic potential analysis reveals a positively-charged exterior in the MACPF β-barrel just above the membrane surface and a negatively-charged interior which may contribute to a charge-repulsion mechanism of lipid clearance on pore formation (Fig 3E). Together with residues from the P2 β-hairpin exterior, the positively-charged exterior patch presumably by contrast helps stabilize its interaction with negatively-charged membrane components such as the lipopolysaccharides found in bacterial cell outer membrane leaflets (Fig 3E).

**Figure 3.**
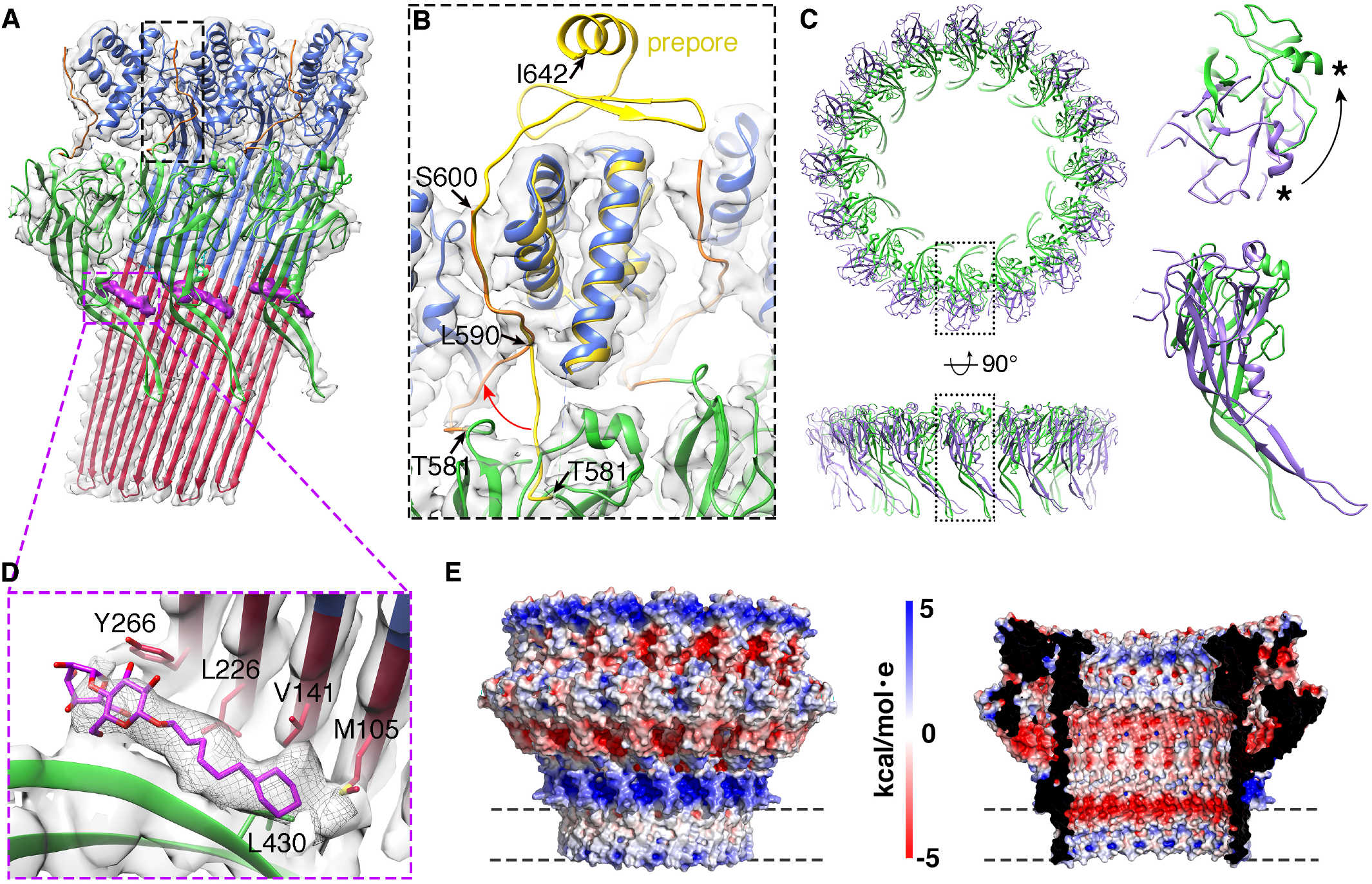
High resolution features of mPFN2 pore. A. Three subunits of mPFN2 pore with a partial CTT (T581-S600, orange) fitted into the density map, boxed with a black rectangle. The bright purple regions show Cymal-6 density identified between the β-barrel and P2 β-hairpins. B. Close-up view of the black box in A, with comparison of the CTT domain between pre-pore (in yellow) and pore (in orange) structures. Pre-pore and pore structures were aligned based on the top helices in the MACPF domain. The red arrow indicates the shift of T581-L590 in the CTT from pre-pore (yellow) to pore (orange). Black arrows point to the highlighted residues. S600-I642 is not resolved in the pore. C.. Overlay of P2 domains from mPFN2 pre-pore ring (dark purple) and pore (green), viewed in two orthogonal orientations. The close-up views of the boxed regions are shown on the right. The P2 domain has a ∼90° rotation between these two states, as indicated by rotation of the same α-helices highlighted with an asterisk. D. Close-up view of cymal-6 binding site. The cymal-6 model (bright purple) is overlapped with its density (gray mesh). The hydrophobic residues surrounding cymal-6 are shown. E. Surface electrostatic potential of the mPFN2 pore-forming oligomer, viewed from outside and inside. The surface electrostatic potential was calculated with APBS plugin in PyMOL. Dash lines indicate the region of detergent belt. The potentials are on a [−5, 5] red-white-blue color map in units of kcal/(mol·e).

### A potential structural conversion from twisted pore to flat pore

Compared with the flat pore, the twisted mPFN2 pore shows a non-perfect helical conformation with an accumulated rise of the β-strands by 63 Å (Fig 4A). The twist of the β-barrel in the twisted mPFN2 pore resembles that of MAC pores, and especially has a similar architecture to the closed form of the MAC pore previously observed (Menny *et al*, 2018) (Fig EV5). However, the direction of the twist is opposite: the twisted mPFN2 is left-handed in the subunit rise direction while MAC pore is right-handed (Fig EV6). The conformations of mPFN2 subunits in the twisted pore are almost identical to those found in the flat pore (pairwise root mean square deviation (RMSD) = 0.085 to 1.204 Å), although the transmembrane β-hairpins in the MACPF domain bend more towards the pore interior (Fig 4B and C). The intersubunit hydrogen-bonding register of the twisted pore remains unchanged (Fig 4B, purple lines), indicating that the accumulated rise is caused by a mild twist of the β-barrel rather than a shift in hydrogen bond register between adjacent subunits (Fig 4B). The plasticity of the transmembrane β-hairpins therefore determines the structural variation in pores, suggesting a relatively straightforward conversion can be made from twisted to flat forms of the pores (Movie EV3); the twisted form may therefore constitute a transition state for pore formation by complete rings of subunits, or alternatively could be an independently-functioning pore-forming state of the PFN2 oligomer.

**Figure 4.**
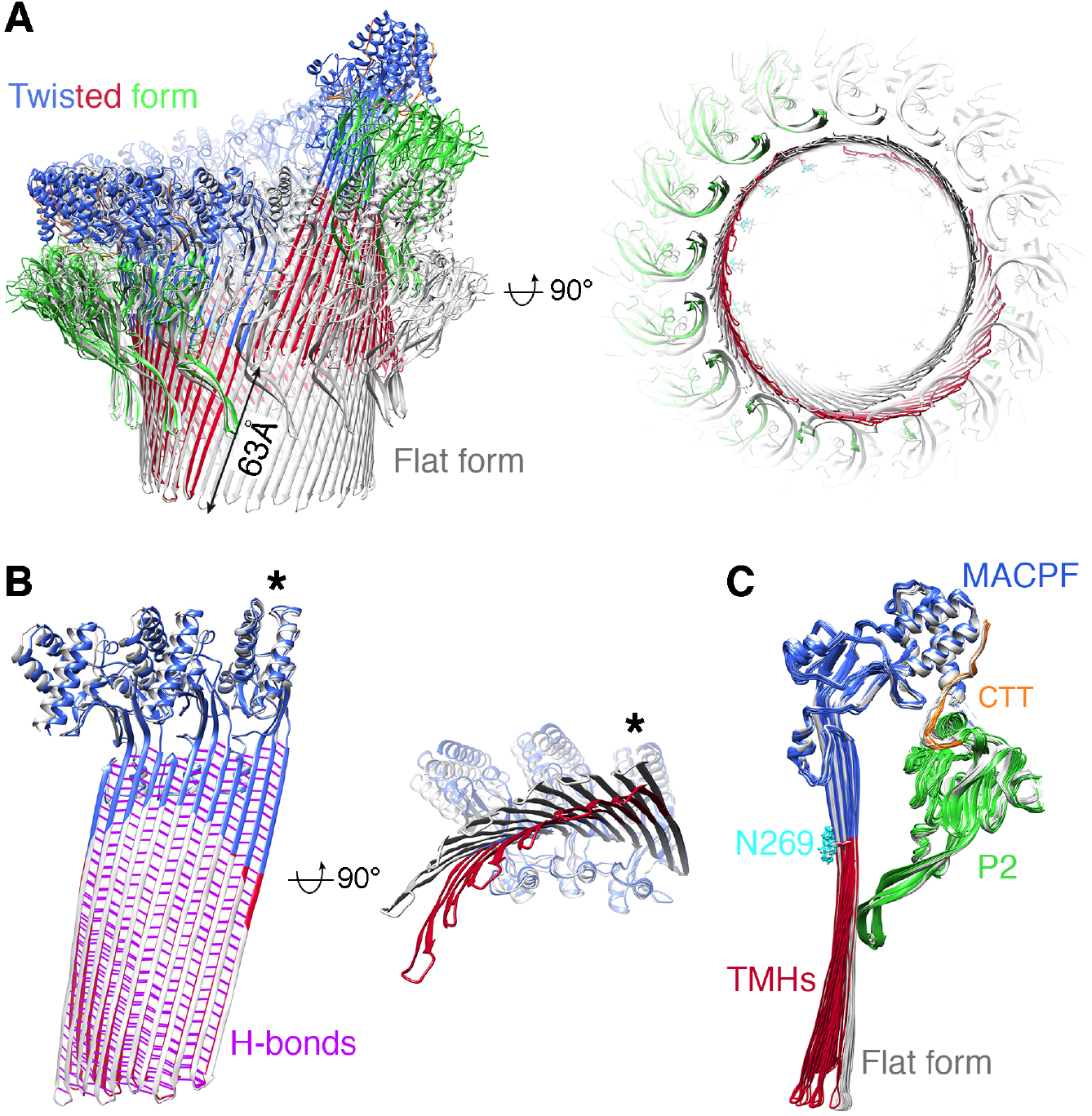
Comparison of the two mPFN2 pore-forming conformations. A. Overlay of twisted pore structure (color-coded as in Fig.1C) with the flat ring pore structure (in gray) viewed from the front and the bottom. The accumulated rise in β-sheet from the twisted form is 63Å as labeled. B. Overlay of three subunits from the twisted pore (colored) with those from the flat ring pore (in gray), shown in two orthogonal views. The hydrogen bonds (H-bonds) in the β-barrel are colored in purple. The asterisk indicates the subunit used for alignment. C. Superposition of all mPFN2 subunits in the twisted (color-coded) and flat conformations (gray), showing an inward curling of the β-stranded transmembrane region in the twisted conformation, RMSD = 0.085 to 1.204Å.

The twisted form also shows a mismatch in the detergent belt, which corresponds to the position of the lipid bilayer with respect to a membrane-inserted form of this oligomer (Fig 2B and E). This mismatch of membrane bilayer position would cause a distortion of the membrane, and we hypothesize that it may play a role in destabilizing the membrane and facilitating pore formation. On the other hand, the membrane bilayer may constrain the twisted form and pull it back into the flat conformation seen for the other pore assembly resolved here in isolation (Fig 2A-C) (which in any case seems the more stable form, as it dominates the population ∼4:1). In either case, the presence of a stable sub-population of pores in a twisted conformation indicates the presence of an alternative state. The oligomeric twist is further manifested in the 3D variability analysis of the twisted pore, which reveals substantial variability of expansion and twisting in pores (Fig EV5B and Movie EV2). These alternative states indicate bases on which mPFN2 might perturb the free energy of the membrane/pore system as a whole, and critically *en route* to pore formation.

### PFN2 pore oligomers formed on membranes are open arc pores

We next attempted to convert the mPFN2 pre-pores formed on a membrane into pores *in situ* by acidification. However, pore formation by mPFN2 at low pH led to aggregation of liposomes, which impedes cryo-ET imaging. Therefore, instead of acidifying the membrane-bound mPFN2 pre-pore, the soluble mPFN2 ectodomain at neutral pH was added to the acidified liposomes (pH 3.6-3.8). Liposomes containing sphingomyelin were used for cryo-ET analysis, which were found to be less prone to aggregation in the presence of mPFN2, due to its reduced binding capacity (Ni *et al*., 2020). Although very sparse on the liposome, mPFN2 pores can be identified confidently from tomograms (Fig 5A). The asymmetry and heterogeneity of the mPFN2 pores (incomplete arc pores) can be observed in side views (Fig 5B). mPFN2 pores do not form junctions between two liposomes, as hPFN2 did (Pang *et al*., 2019); but such packing of pore complexes head-to-head is a well-known artefact of some cryo-EM analyses of pore complexes (Gilbert, 2021).

**Figure 5.**
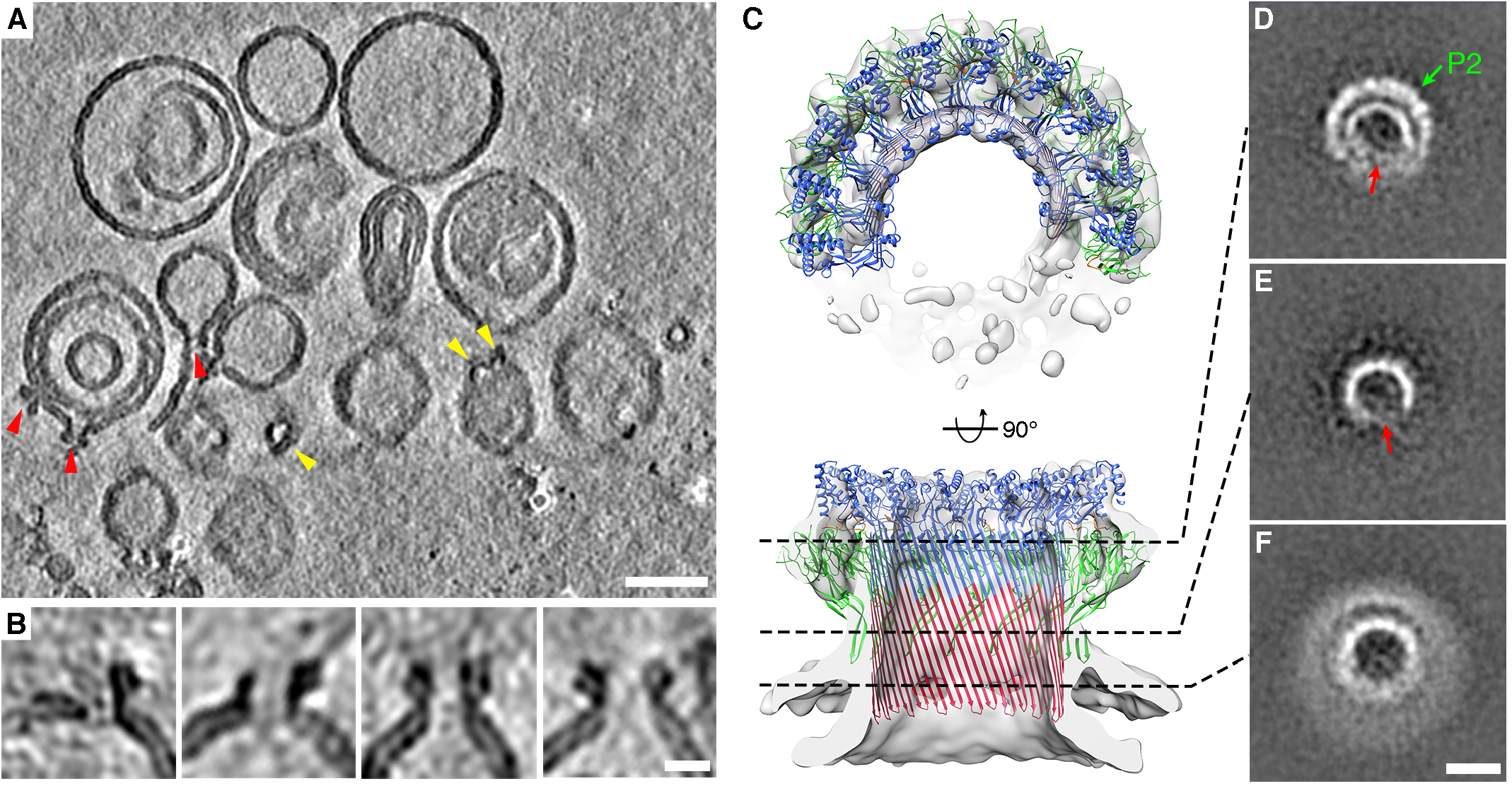
CryoET and subtomogram averaging of mPFN2 pores on sphingomyelin liposomes. A. A representative tomogram slice of a mPFN2 pore on a membrane. Red and yellow arrowheads point to pores on membrane with side and top views, respectively. Slice thickness 32.6 Å. Scale bar = 50 nm. B. Close-up examples of mPFN2 pores on a membrane shown as side views, highlighting the asymmetry and variability of pore structures. Scale bar = 10 nm. C. Subtomogram averaging of mPFN2 pores on liposomes, at 18 Å resolution. The arc-form pore is shown as a top view (upper panel) and a central slice through a side view (lower panel). D-F. 2D cross-section slices along the pore at different heights. The green arrow points to the P2 domain in the periphery and red arrows point to the opening of the arc-pore. Scale bar = 10 nm.

Subtomogram averaging of the mPFN2 pores *in situ* on membranes revealed an incomplete arc pore structure at a resolution of ∼18 Å (Fig 5C-F and Fig EV2B), reflecting the heterogeneity and openness of pores on membranes. Since most of the pre-pore oligomers in isolation and on the membrane are arcs at neutral pH (Fig EV1A-B and Fig1A-B), we strongly suspect that the arcs of subunits in a pore-forming state we observe in the membrane context have only reached partial ring assembly (pre-pore arcs) before pore formation occurs *in situ*. The curvature of the β-barrel from the arc pore on the membrane is similar to that of the ring pore in isolation, as indicated from a rigid-body fitting of a partial pore structure with ten subunits into the density map (Fig 5C). Importantly, the pore structures on liposomes confirm that the P2 domain bound to a targeted membrane adopts the same conformation as that observed in the isolated pore, i.e., the tip of the β-hairpin points downwards to interact with the same membrane in which the MACPF domain forms the pore.

These structures of mPFN2 at different states and conformations enable us to propose the following mechanism of PFN2 transition. PFN2 forms pre-pore arcs and rings on membranes at neutral to slightly acidic pH conditions (Fig 6A) and converts to pores upon acidification. Due to the opposing orientation of the MACPF domain and its target membrane in the pre-pore form, for pre-pores with complete rings of subunits a transitional opening is required as an intermediate step to avoid the steric clash of a rotating MACPF domain against its neighbors during transition. The twisted pore structure provides such an opened form/transition state and we propose the following steps in pre-pore to pore conversion for a complete pre-pore ring of subunits: pH-induced cracking and twisting of the oligomeric pre-pore ring at one critical point allows rotation of the MACPF domain (Fig 6B, Movie EV4); the MACPF domain from the first subunit at the break point flips 180° (Fig 6C); and then refolds and inserts into the membrane (Fig 6D). The twisted pore structure may thus act as an intermediate pore stage (Fig 6E), or quickly convert to a flat ring pore structure on the membrane, perhaps influenced by its interaction with the membrane bilayer (Fig 6F). In the case of an incomplete pre-pore ring (an arc of subunits), the twisting breakage is not necessary as the pH-induced rotation of the first MACPF domain to transition to a pore state will already be sterically allowed at its open end, but can propagate throughout the rest of an arciform pre-pore oligomer in the same way as would occur for the twisted pore. In any case, steric constraints require that the pre-pore to pore transition propagates among subunits.

**Figure 6.**
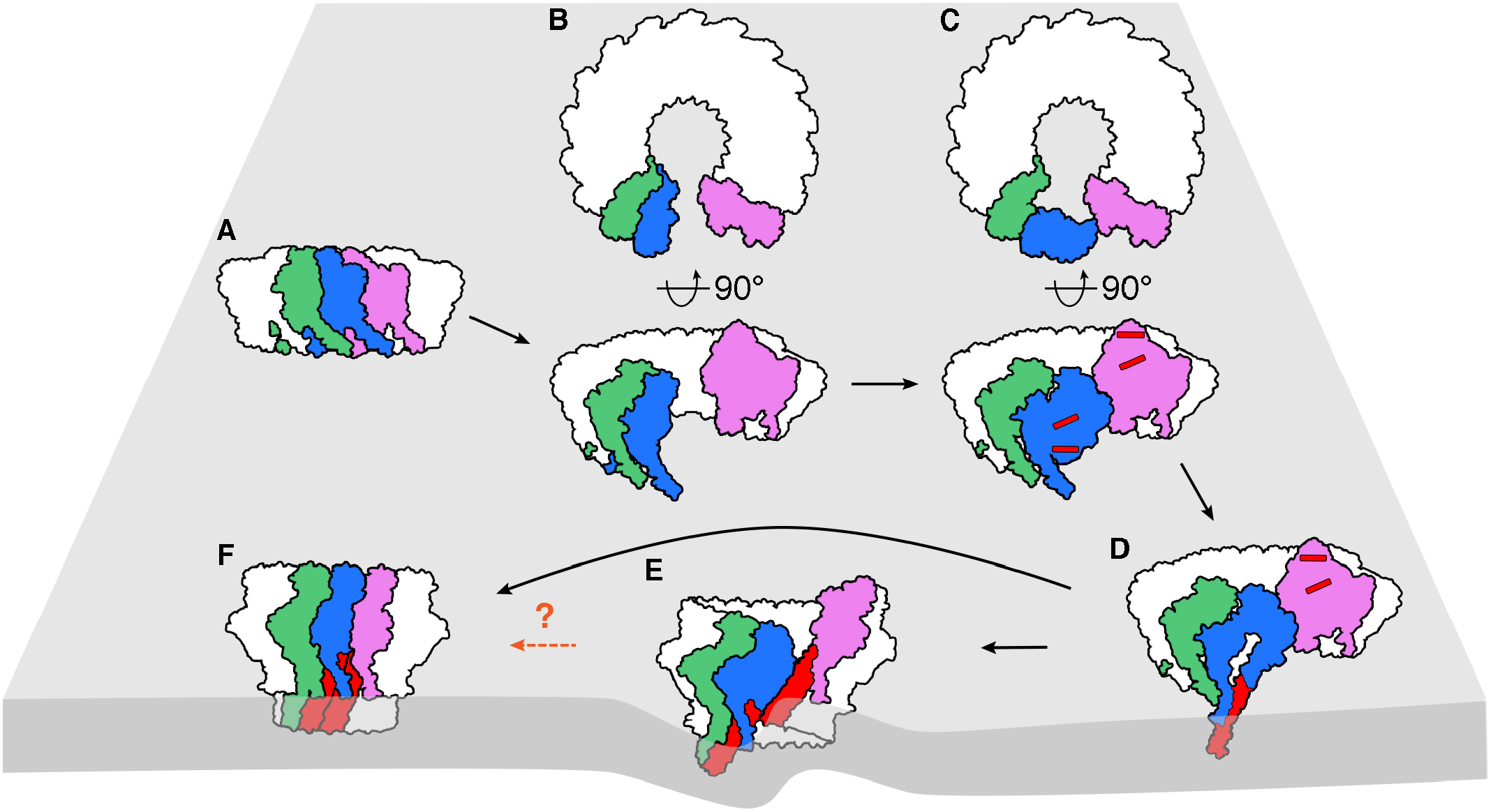
Proposed model of mPFN2 pre-pore to pore conformational change. A. Ring of subunits in a pre-pore conformation bound to a membrane surface, anchored at the ring subunits’ P2 domain. Three subunits of the ring are colored in green, blue and violet. B. A twisted pre-pore state forms in two orthogonal views. C. The MACPF domain of one subunit (blue) at the cracking point undergoes a ∼180° rotation so that the transmembrane-hairpin helices in the pre-pore subunit (red bars) face the membrane to which the P2 domain is bound. D. The transmembrane hairpins (red) of the blue subunit are deployed to penetrate the targeted membrane. E. A second subunit (green) can now undergo the same transition as the blue subunit, and the conformational change propagates towards the other end of the cracking point. A twisted pore is then formed, which distorts the membrane. F. The twisted pore could ultimately convert to a flat ring conformation on the membrane or remain twisted. The dashed arrow and question mark indicate that the twisted form could be an intermediate state or a final state.

## Discussion

Perforin-2 is the most ancient MACPF family protein characterized in animals and has been implicated in multiple biological pathways. Together with our previous work (Ni *et al*., 2020), we have determined structures of the mPFN2 pre-pore and pore in isolation and on a membrane, including *via in situ* assembly. These snapshots have revealed a mechanism of pore formation by mPFN2 which enables large conformational changes to be achieved, firmly gating the transition from pre-pore to functional pore. mPFN2 will form small oligomers in solution but on binding to a membrane generates pre-pore assemblies with varying degrees of ring completeness (Fig 1 and Fig EV1). mPFN2 can also oligomerize into a full-ring pre-pore in isolation, at a moderately low pH (∼pH 5.5) (Ni *et al*., 2020) and these pre-pore rings can transit into both flat and twisted pores, with the same number of subunits (Figs 2 and 4). The higher resolution achieved here for both flat and twisted pore conformers enables a detailed dissection of conformational changes associated with the pre-pore to pore transition, while anchored by the P2 domain and especially its β-hairpin tip on a targeted membrane in which pore formation occurs (Figs 2 and 5). Pores formed on a membrane are predominately arcs instead of complete rings of 16 subunits (whether flat or twisted), but all show a similar domain organization and curvature to pore structures formed in isolation (Fig 5). Together, these results suggest that the MACPF domain rotates 180° with respect to the membrane-bound P2 domain during pre-pore to pore transition on membranes, with P2 domains remaining attached to the target membrane via their extended β-hairpins (Fig 6).

The different oligomerization states of mPFN2 oligomers reveal that constraints on its self-association control the precise conformation of a MACPF/CDC assembly. These constraints include whether it is bound to a membrane or free in solution and undergoing a concentration-driven self-association. For example, when an appropriate membrane support (such as a negatively charged lipid bilayer or liposome) is available, mPFN2 oligomerizes efficiently to form a ring or arc pre-pore even at a neutral pH, whereas in solution, mPFN2 forms small oligomers in the majority of cases (Fig EV1). This is as expected for proteins whose oligomerization and pore formation are driven by a kinetic mechanism (Gilbert, 2005; Gilbert *et al*, 2014; Gilbert & Sonnen, 2016; Leung *et al*, 2014). After converting to a pore-forming structure in isolation, without membrane restraints, 16-meric assemblies that are both flat and ring-shaped and twisted structures may be observed (Fig 2) (Ni *et al*., 2020). However, on the membrane, where diffusion is more restricted and local depletion of free monomer is more likely (Gilbert, 2005), the majority of the pre-pores would not have grown into complete rings and the resulting pores are more likely to be in the open arc form, as we observe (it is also possible that pore formation by arcs of subunits is less dependent on acidification than pore formation by complete rings). These toroidal pores are similar to the previous observation of incomplete pneumolysin pores (Gilbert & Sonnen, 2016; Sonnen *et al*, 2014), and also to suilysin (Leung *et al*., 2014), listeriolysin (Podobnik *et al*, 2015; Ruan *et al*, 2016), MAC complex (Sharp *et al*, 2016) and gasdermin family proteins (Mulvihill *et al*, 2018) on membranes, which display open arcs or “slit” pores. These observations are entirely in accord with the model of pore formation by kinetically trapped MACPF/CDC oligomers previously articulated (Gilbert, 2005; Gilbert *et al*., 2014; Gilbert & Sonnen, 2016; Leung *et al*., 2014). Our work confirms the capacity of MACPF/CDC proteins – and here, specifically mPFN2 – to form pores using incomplete arcs of subunits (Fig 5) as well as complete rings (Figs 2 and 4). We believe that the adaptively preferred biological outcome may be a pore made of an incomplete ring of subunits and complemented by a hemi-toroidal lipid bilayer structure, in which scenario less pore-forming reagent is required for functional pore formation. It is also possible that the twisted pore state we observe for mPFN2 is of relevance to the pre-pore to pore conversion of full rings of subunits with MACPF/CDC proteins too: as well as the MAC itself helical arrangements of subunits have been observed for the CDC pneumolysin (Gilbert *et al*, 1999) and the *Toxoplasma gondii* perforin-like protein 1 (Ni *et al*, 2018).

Our study focused on the mechanism of pore formation by soluble mPFN2 as would occur following its cleavage from its transmembrane anchor (McCormack & Podack, 2015; McCormack *et al*., 2015a). It is possible that the delivery of IL-33 or some other aspect(s) of its biological activity involve pore formation by mPFN2 when it is still membrane anchored (without being cleaved off the membrane), which merits further investigation.

## Materials and Methods

### Protein expression and purification

Ectodomain of mPFN2 (Uniprot: A1L314, residues 20-652) was expressed in HEK293T cells and purified from the media using anti-1D4 antibody-coupled agarose resin as previously described (Ni *et al*., 2020). The proteins were concentrated to ∼1.5 mg/ml in 20 mM Hepes (pH7.5) and 300 mM NaCl, before flash-frozen in liquid nitrogen with 10 µL aliquots. The protein aliquots were saved in −80 °C freezer until further use.

### Cryo-EM SPA sample preparation and data collection

Cryo-EM grids of mPFN2 pre-pore oligomers at pH7.5 were prepared by directly freezing the purified protein at ∼ 1.5 mg/ml onto holey lacey-carbon grids (Agar Scientific, UK) using a Vitrobot Mark IV (Thermo Fisher Scientific). The grids were then imaged using a Thermo Scientific Glacios equipped with a Falcon 3 camera operated in linear mode. The mPFN2 pore sample was prepared as described previously (Ni *et al*., 2020). In brief, purified mPFN2 protein was firstly assembled as pre-pore oligomers by incubating with buffer containing 50 mM Citric acid (pH 5.5) and 150 mM NaCl at 0.5 mg/ml at 37°C overnight. The pre-pore structures were then converted to pore structures by lowering pH to 3.6∼4.0 in the presence of 0.056% Cymal-6 at room temperature for 30 min. To prepare cryo-EM grids, 3 μl of mPFN2 pore (0.2 mg/ml) was applied to a glow-discharged lacey-carbon grid with ultra-thin carbon support film (Agar Scientific, UK), blotted for 2.5 s in 100% humidity at 22 °C and plunged into liquid ethane using a Vitrobot Mark IV (Thermo Fisher Scientific). The mPFN2 pore dataset were acquired using a Thermo Scientific Krios electron microscope operating at 300 kV via EPU, with a Gatan K2 Summit direct electron detection camera operated in counting mode.

### Cryo-EM single particle analysis

The pre-pore dataset was processed in cryoSPARC v3.3.2 (Punjani *et al*, 2017), following the standard pipeline including micrographs motion correction and CTF estimation. The particles were initially picked though blob picker. The initial 2D class averages revealed clear separation of mPFN2 assemblies, and were subsequently used as templates for template-based picking. Several rounds of 2D classification were performed to remove junk particles. The particles display strongly preferred orientations, as all the 2D class averages are in top views, which impedes further processing like 3D classification and refinement. Cryo-EM SPA of isolated mPFN2 pore structures were conducted using RELION (Scheres, 2012) and the procedure is outlined in Figure EV3. Briefly, beam-induced motion was corrected with MotionCor2 (Zheng *et al*, 2017) to generate dose-weighted micrographs from all frames. The contrast transfer function was estimated using Gctf (Rohou & Grigorieff, 2015; Zhang, 2016). A total of 1,143,742 particles were picked using LoG-based autopicking from RELION and further cleaned by several rounds of 2D classification. 3D classification in RELION with C1 symmetry resulted in two major classes: the flat ring form and twisted form. The flat ring form maps were refined using C1, C2 and C16 symmetries, and application of C2 symmetry resulted in the best quality map for both MACPF and P2 domains at ∼3 Å. Refinement with C16 symmetry improved the overall resolution to 2.7Å, but density of the peripheral P2 domain was less resolved. Furthermore, the bottom of the MACPF β-barrel is slightly elliptical in the C1 and C2 map, indicating a deviation from C16 symmetry in the structure. The twisted form was refined with C1 symmetry at an overall resolution of ∼4 Å. The overall B factor was determined by the RELION PostProcess tool. Resolution of the maps was determined by the gold-standard Fourier shell correlation (FSC) at 0.143 criterion, with a soft mask applied around the protein density. The local resolution was calculated using RELION and presented in Chimera.

### 3D variability analysis of cryo-EM SPA

The flat and twisted forms of mPFN2 pores were classified in RELION by 3D classification with C1 symmetry, and imported to cryoSPARC (Punjani *et al*., 2017) for 3D variability analysis, respectively. The first two modes were solved for both classes and 20 consecutive conformational maps were generated for both forms.

### Model building, refinement and structure visualization

The pre-pore structure in solution (PDB accession code 6SB3) was rigid-body fit into the cryoET subtomogram averaged density and the β-hairpin in the P2 domain was fit into the density manually before further refinement using molecular dynamics flexible fitting with Namdinator (Kidmose *et al*, 2019). For both flat ring and twisted pore maps from cryo-EM SPA, the previous pore structure (PDB accession code 6SB5) was used as the initial model, with further real-space refinement in Coot (Emsley & Cowtan, 2004) and phenix.real_space_refine (Afonine *et al*, 2012). The figures were prepared using Chimera (Pettersen *et al*, 2004) and PyMOL (The PyMOL Molecular Graphics System, Version 2.0 Schrödinger, LLC).

### Cryo-ET sample preparation and data collection

To prepare mPFN2 prepore oligomers on membranes, liposomes composed of *E*.*coli* total lipid extracts (Avanti Polar Lipids Inc) were prepared at a concentration of 10mg/ml in 20 mM HEPES (pH 7.5) and 150 mM NaCl, and extruded through a 50 nm polycarbonate membrane (Whatman). mPFN2 protein at a final concentration of 0.6 mg/ml was added to the freshly prepared liposomes (final liposome concentration: 3 mg/ml) and incubated at 37°C for 1h. The protein-liposome mixture was mixed with 6 nm gold fiducial markers and 3 ul was applied to the carbon side of glow-discharged lacey carbon film grid (AGS166-3, Agar Scientific). The grids were blotted for 3∼4 s at 100% humidity at 22°C and flash frozen in liquid ethane using a Vitrobot Mark IV (Thermo Fisher Scientific).

To prepare mPFN2 pore oligomers on liposomes at low pH, liposomes with a composition of POPC/SM (1-palmitoyl-2-oleoyl-sn-glycero-3-phosphocholine and sphingomyelin, (brain, Porcine) at a 1:1 (w:w) ratio) were prepared at a concentration of 10 mg/ml in 50 mM citric (pH3.6-3.8) and 150 mM NaCl, extruded through a 50 nm polycarbonate filter (Whatman). Liposomes at a final concentration of 2mg/ml and mPFN2 at a final concentration of 0.5mg/ml were mixed and incubated at 37°C for 1h. The cryoET grids were prepared in the same manner as describe above.

Tilt-series were collected on a 300 kV Krios microscope (Thermo Fisher Scientific) equipped with electron counting cameras using serialEM (Mastronarde, 2005) and TOMO. A dose-symmetric data collection scheme (Hagen *et al*, 2017) was used, with a tilt angle ranging from −60° to 60° at a 3° step and group of 3. A total electron exposure of ∼123 e^-^/Å^2^ were applied to each tilt-series. The details of data collection parameters are summarized in Supplementary Table EV1.

### Cryo-ET subtomogram averaging

Cryo-ET subtomogram averaging was performed using emClarity (v1.5.3.10 (Himes & Zhang, 2018; Ni *et al*, 2022)) and Dynamo (Castano-Diez *et al*, 2012). For the pre-pore on E.coli liposomes dataset, an initial map from cryo-EM SPA (EMD-10134) was used as a template to pick the pre-pores in emClarity. CTF multiplied tomograms were binned at 6× (hereafter bin6 tomograms) and the template was filtered at a resolution ∼30 Å determined automatically inside emClarity. A total of 25,530 subtomograms were picked and the false positives were initially removed by 3D classification after a few iterations of alignment at bin4. The resulting dataset (17,180 subtomograms) were then checked manually to remove the “junk particles”, which were mainly membrane bilayers and isolated pre-pores in solution. The final dataset containing 11,794 subtomograms was imported to Relion4 and refined using C1 symmetry with bin2 pseudo-subtomograms, before 3D classification with a molecular mask applied on the mPFN2 pre-pore. The class which showed a complete ring stage pre-pore was selected (3,695 subtomograms) and further refined using C16 symmetry at bin2, with a molecular mask applied excluding membrane density during refinement. Per-particle motion was estimated in Tomo frame alignment and final reconstruction was performed from 2D tilt-series and postprocessed.

mPFN2 pores bound to sphingomyelin liposomes were identified manually in IMOD (Mastronarde & Held, 2017), using the SIRT-like filter tomogram reconstructions at bin8. A total of 1,369 pores were identified from 141 tomograms. The subtomograms were initially extracted from the bin8 tomograms above and aligned with a reference map generated *ab initio* in Dynamo (Castano-Diez *et al*., 2012). The orientation and coordinate of the refined subtomograms were manually checked and only the ones with correct orientation and position were kept for further alignment. The resulting dataset containing 992 subtomograms were imported to emClarity, and the subsequent alignment and averaging steps were performed in emClarity using 6×, 4× and 2× binned tomograms, respectively. The final two half density maps were reconstructed using cisTEM within the emClarity package and combined using Relion postprocess tool.

## Acknowledgments

We thank Yanan Zhu and Yun Song for technical support in cryo-EM and ET data acquisition. We acknowledge Diamond Light Source for access and support of the cryo-EM facilities at the UK national electron bio-imaging center (eBIC, proposal EM20223 and NT21004), funded by the Wellcome Trust, MRC, and BBSRC. The Division of Structural Biology is a part of the Wellcome Centre for Human Genetics, Wellcome Trust Core Grant Number 090532/Z/09/Z. Electron microscopy provision was provided through the OPIC electron microscopy facility, a UK Instruct-ERIC Centre, which was founded by a Wellcome JIF award (060208/Z/00/Z) and is supported by a Wellcome equipment grant (093305/Z/10/Z). Computation was performed at the Oxford Biomedical Research Computing (BMRC) facility, a joint development between the Wellcome Centre for Human Genetics (Wellcome Trust Core Award Grant Number 203141/Z/16/Z) and the Big Data Institute (BDI) supported by Health Data Research UK and the NIHR Oxford Biomedical Research Centre. X.Y. and R.J.C.G. were supported by the Calleva Research Centre for Evolution and Human Sciences at Magdalen College, Oxford. T.N. and P.Z. were supported by the UK Wellcome Trust Investigator Award 206422/Z/17/Z, the UK Biotechnology and Biological Sciences Research Council grant BB/S003339/1, and the ERC AdG grant (101021133).

## Author Contributions

X.Y.: Conceptualization; Data curation; Formal Analysis; Investigation; Methodology; Validation; Visualization; Writing – original draft; Writing – review & editing

T.N.: Conceptualization; Data curation; Formal Analysis; Investigation; Methodology; Project administration; Software; Supervision; Validation; Visualization; Writing – original draft; Writing – review & editing

G.M.: Conceptualization; Writing – original draft; Writing – review & editing

P.Z.: Funding acquisition; Methodology; Supervision; Software; Writing – original draft; Writing – review & editing

R.J.C.G.: Conceptualization; Formal Analysis; Funding acquisition; Investigation; Methodology; Project administration; Resources; Supervision; Visualization; Writing – original draft; Writing – review & editing

## Data availability

Electron microscopy and subtomogram averaging maps and structures reported in this paper are being deposited in the RCSB PDB and in the Electron Microscopy Data Bank. Accession codes: STA of mPFN2 pre-pore and pore on membrane EMD-15076 and EMD-15087; SPA of mPFN2 pore flat form EMD-15072, PDB 8A1D; twisted form EMD-15086, PDB 8A1S.

## Figure legends

**Figure EV1**. mPFN2 oligomers in solution at pH7.5

A. Representative 2D class averages of mPFN2 assemblies in solution at pH7.5 (pre-pore state). Number of subunits for each species are labelled.

B. Distribution of assemblies forming in solution from data showcased in A, from one to multiple subunits.

**Figure EV2**. Gold standard half map FSC of mPFN2 pre-pore and pore from subtomogram averaging.

A. FSC of mPFN2 pre-pore on membrane.

B. FSC of mPFN2 pore on membrane.

C. Local resolution of mPFN2 pre-pore estimated in Relion.

D. Central slice of mPFN2 pre-pore map in a sideview showing smearing of membrane density under mPFN2 pre-pore. Scale bar = 10 nm.

**Figure EV3. Cryo-EM data processing workflow for mPFN2 pore structures in isolation**.

The diagram illustrates the image processing and 3D reconstructions for mPFN2 pore in different conformations. Maps from the C2 symmetry in the flat ring conformation were used to generate the final map because the bottom of the β-barrel does not follow C16 symmetry.

**Figure EV4. Cryo-EM single particle analysis of mPFN2 pore structures in isolation**.

A. Gold-standard FSC curves of flat ring pore maps refined with C1, C2 and C16 symmetries and the twisted pore map with C1 symmetry. Soft masks were applied to two half-maps in relion postprocessing.

B. Model-to-Map FSC curves showing the correlations between the refined atomic models and their corresponding cryo-EM maps.

C. Local resolution of the mPFN2 pore in the ring (upper) and twisted (lower) conformations.

D. Representative regions of cryo-EM densities (gray mesh) superposed with an atomic model of the mPFN2 pore in ring conformation and colored as in Fig.1E.

**Figure EV5**. 3D variation analysis of mPFN2 pore structures in isolation. Twenty classes were generated for both flat and twisted pores.

A. Two representative 3D classes of mPFN2 ring conformations showing the β-barrel varies between circular and elliptical forms. Black dashed boxes indicate the consistency of circularity in the top of MACPF domain, while the orange dashed boxes show the variability in the bottom of the MACPF domain.

B. The 1^st^ and 20^th^ 3D classes of mPFN2 twisted forms viewed from the top and the side, showing the variability of twist.

**Figure EV6**. Comparison of MAC and mPFN2 pores.

A-B Structures of MAC in two different conformations (PDB: 6H03 and 6H04).

C Structure of mPFN2 in the twisted pore conformation. The direction of twist is indicated by the arrows.

