## Supplementary material for "Structures of perforin-2 in solution and on a membrane reveal mechanisms for pore formation": EV Supplementary Information

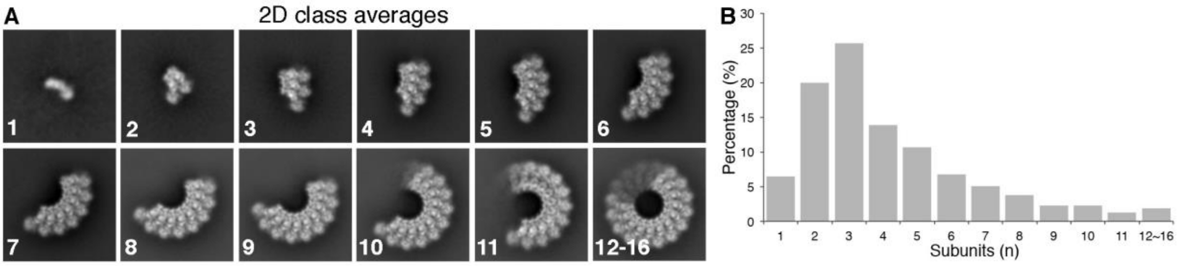


**Figure EV1**. mPFN2 oligomers in solution at pH7.5

A. Representative 2D class averages of mPFN2 assemblies in solution at pH7.5 (pre-pore state). Number of subunits for each species are labelled.
B. Distribution of assemblies forming in solution from data showcased in A, from one to multiple subunits.

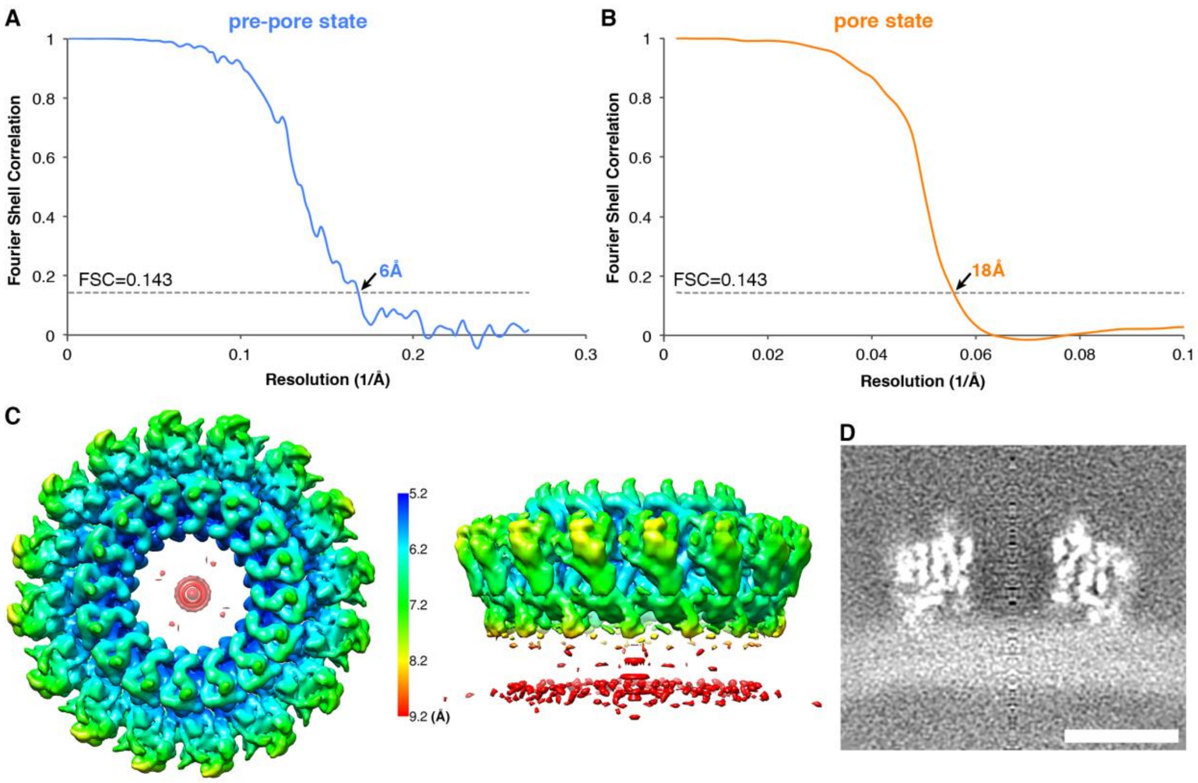


**Figure EV2**. Gold standard half map FSC of mPFN2 pre-pore and pore from subtomogram averaging.

A. FSC of mPFN2 pre-pore on membrane.
B. FSC of mPFN2 pore on membrane.
C. Local resolution of mPFN2 pre-pore estimated in Relion.
D. Central slice of mPFN2 pre-pore map in a sideview showing smearing of membrane density under mPFN2 pre-pore. Scale bar = 10 nm.

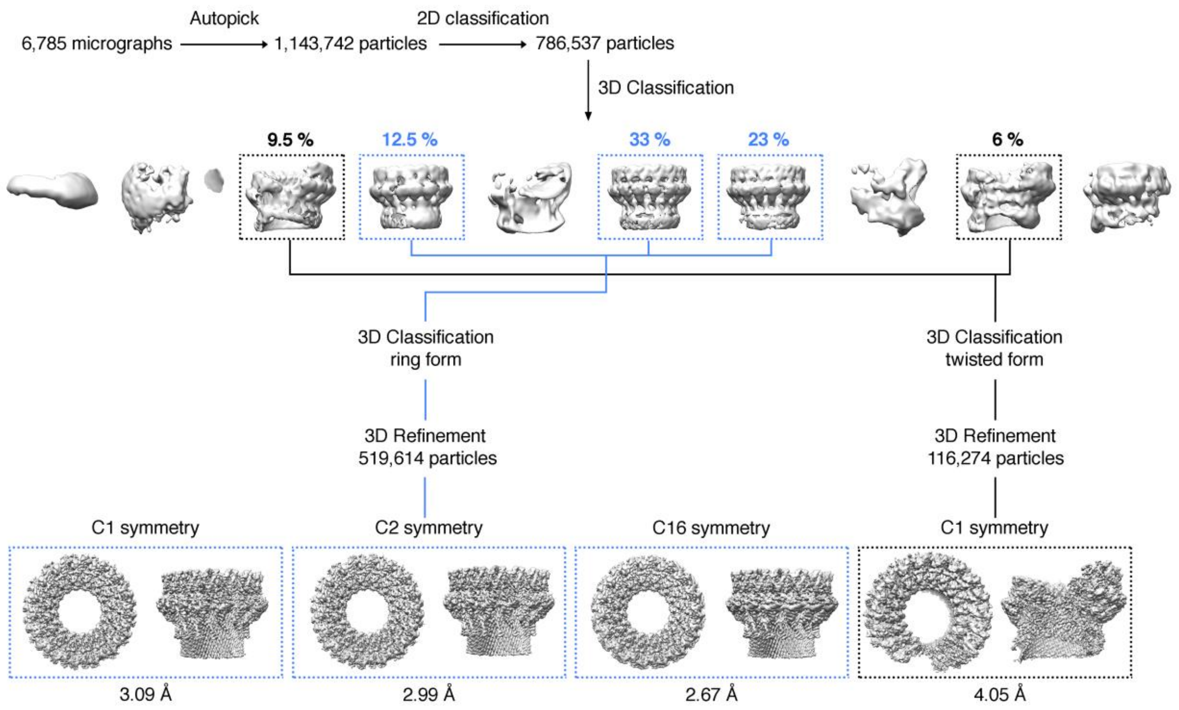


**Figure EV3. Cryo-EM data processing workflow for mPFN2 pore structures in isolation.**

The diagram illustrates the image processing and 3D reconstructions for mPFN2 pore in different conformations. Maps from the C2 symmetry in the flat ring conformation were used to generate the final map because the bottom of the β-barrel does not follow C16 symmetry.

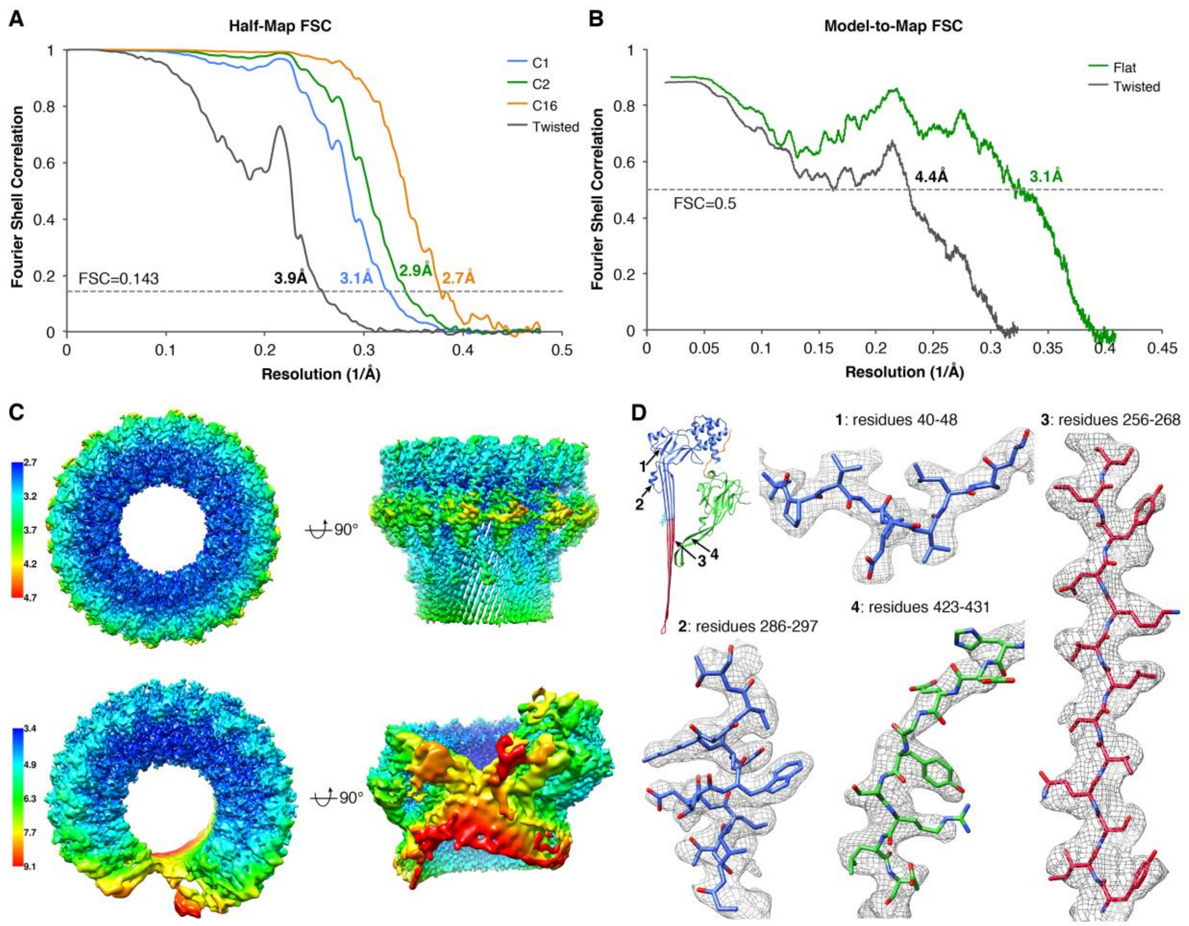


**Figure EV4. Cryo-EM single particle analysis of mPFN2 pore structures in isolation.**

A. Gold-standard FSC curves of flat ring pore maps refined with C1, C2 and C16 symmetries and the twisted pore map with C1 symmetry. Soft masks were applied to two half-maps in relion postprocessing.
B. Model-to-Map FSC curves showing the correlations between the refined atomic models and their corresponding cryo-EM maps.
C. Local resolution of the mPFN2 pore in the ring (upper) and twisted (lower) conformations.
D. Representative regions of cryo-EM densities (gray mesh) superposed with an atomic model of the mPFN2 pore in ring conformation and colored as in Fig.1E.

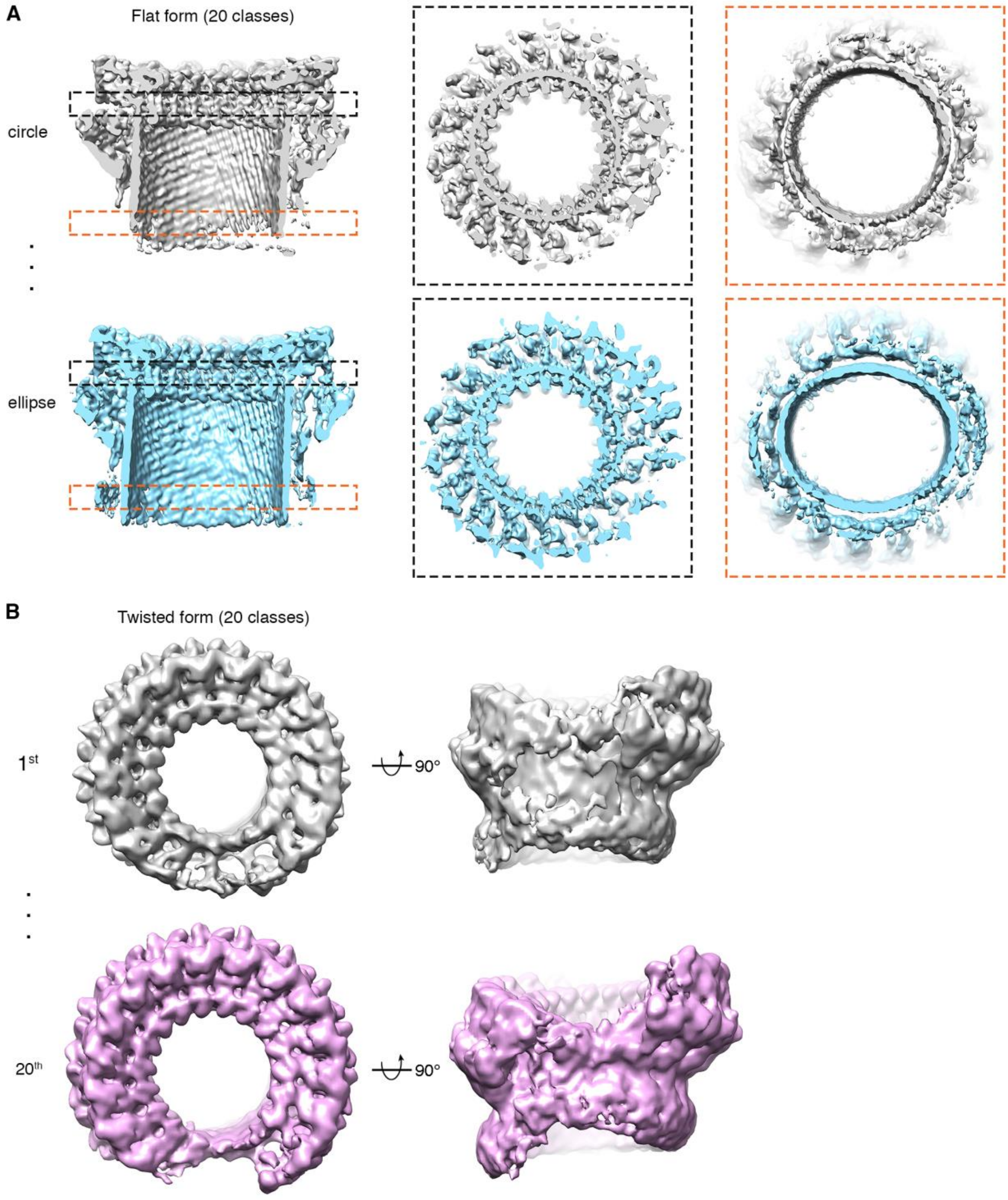


**Figure EV5.** 3D variation analysis of mPFN2 pore structures in isolation. Twenty classes were generated for both flat and twisted pores.

A. Two representative 3D classes of mPFN2 ring conformations showing the β-barrel varies between circular and elliptical forms. Black dashed boxes indicate the consistency of circularity in the top of MACPF domain, while the orange dashed boxes show the variability in the bottom of the MACPF domain.
B. The 1st and 20th 3D classes of mPFN2 twisted forms viewed from the top and the side, showing the variability of twist.

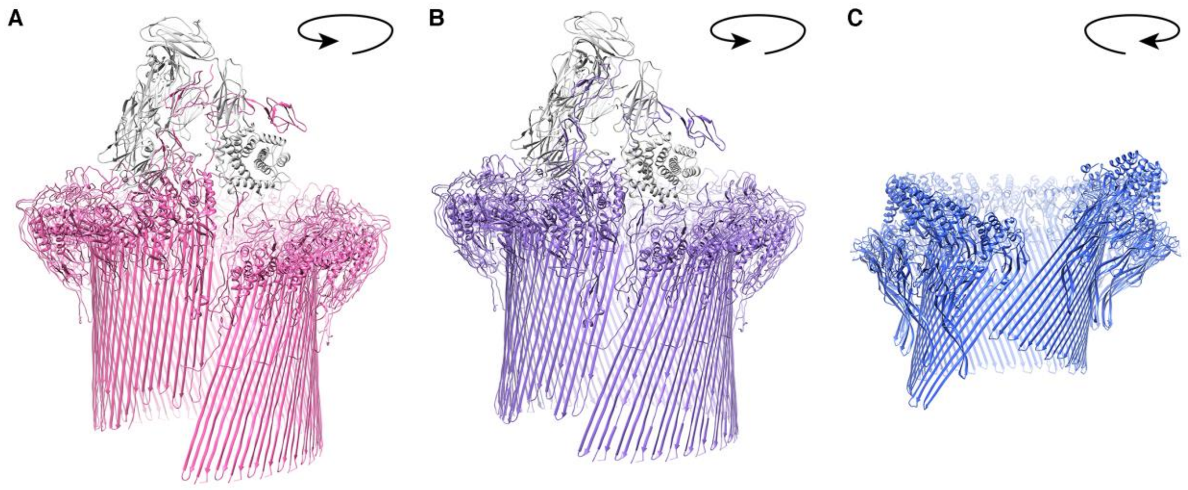


**Figure EV6.** Comparison of MAC and mPFN2 pores.

A-B Structures of MAC in two different conformations (PDB: 6H03 and 6H04).

C Structure of mPFN2 in the twisted pore conformation. The direction of twist is indicated by the arrows.
